# Ripple-Mapping for the Detection of Long Duration Action Potential Areas in Patients with Brugada Syndrome

**DOI:** 10.1101/263145

**Authors:** Rui Providencia, Diogo Cavaco, Pedro Carmo, Cristiano Pisani, Francisco Moscoso Costa, Mariana Faustino, Francisco Morgado, Pier D Lambiase, Mauricio Scanavacca, Pedro Adragao

## Abstract

**Purpose:** Catheter ablation has been recently utilised in patients with Brugada syndrome to prevent ventricular arrhythmias. We hypothesized whether a mapping algorithm “Ripple-mapping” would be able to rapidly identify the areas of long-duration multicomponent electrograms which constitute the targets for ablation for an automated strategy.

**Methods:** Ripple-Mapping analysis was performed in all patients undergoing catheter ablation of Brugada syndrome in 2 centers. The activity propagation pattern determined by Ripple-Mapping was assessed, and areas of long duration potentials identified by visual inspection. The area of interest with long-duration potentials was correlated with the location of the ablation lesions.

**Results:** Detection of long duration potentials was possible in all four patients in this analysis. Points marking the ablation area in 2 patients were a perfect match to abnormal areas identified by Ripple-mapping. However, in 2 patients Ripple-mapping identified additional sites of long duration potentials which were not ablated. Acute and mid-term normalization of the ECG pattern was observed in three patients, and all these were free from arrhythmia relapse during a 10.3±3.2 months follow-up. In one patient, who had no ECG normalization and had arrhythmia relapse, the ablated area corresponded to <25% of the Ripple-map region of interest.

**Conclusions:** This study shows the potential use of Ripple-mapping for identifying areas of long duration multicomponent electrograms in patients with Brugada syndrome and recurrent ventricular arrhythmias. This technique enables the rapid definition of an area of interest which should then be validated by the operator before performing ablation with an end-point of ECG normalisation.

## Background

Brugada syndrome can lead to ventricular arrhythmias and sudden cardiac death [1]. Its underlying pathophysiological mechanism (repolarization vs. depolarization defect) is still unclear [2]. The implantable cardioverter defibrillator is a life-saving therapy and has been the main treatment to protect patients at risk of ventricular fibrillation [3]. However, some individuals experience repeated episodes of ventricular fibrillation and appropriate shocks [4-5].

Catheter ablation has been recently proposed as a therapeutic option [6], in the *European Society of Cardiology* guidelines (a class IIb indication, level of evidence C) for patients with history of electrical storm or multiple appropriate therapies [7]. During this procedure, multicomponent long-duration electrograms are ablated in the epicardial right ventricular outflow tract [6, 8-9]. This involves operator analysis of every single point or, as an alternative, using an automated algorithm for creating potential duration maps [10], which is not currently commercially available.

Conventional mapping of arrhythmias assigns a single value of timing or voltage to each coordinate. This approach may under-represent important areas, such as long amplitude fractionated potentials or double potentials. Furthermore, conventional mapping techniques require precise annotation, with the operator carefully selecting the timing of local activation. This can be challenging with some electrograms-it is highly dependent on operator skill and experience, and if performed inaccurately can invalidate the entire map. Ripple-mapping is a method of electrogram visualization which displays entire electrogram voltage-time relationship at each location, doesn’t’require operator-determined window of interest, manual verification of local activation times, or data interpolation into regions of unmapped myocardium [11]. It has already been used to map atrial tachycardias [12], persistent atrial fibrillation [13], and visualize and target slow conduction channels within infarct-related ventricular scar [14-15].

The precise location and extent of the target substrate in the right ventricle outflow tract & other areas of the right ventricle [6] in patients with Brugada syndrome is highly variable, and detailed mapping and identification through the conventional approach may be a protracted process. Also, areas of interest may be easily missed if the catheter is moved to quickly across a region. We hypothesize that Ripple mapping would standardise & optimise identification of areas of long-duration multicomponent electrograms constituting targets for ablation in Brugada syndrome patients and provide further insight into its pathophysiological mechanism.

## Methods

From April 2016 to March 2017, all patients undergoing catheter ablation in the setting of Brugada syndrome using the CARTO mapping system in two high-volume Cardiac Electrophysiology Centers (Lisboa and São Paulo) were considered eligible for this analysis. The ablation procedures and use of the data for research was approved by the institutional board and local ethics committees. Patients provided informed consent.

### Catheter Ablation Procedure

Procedures were performed under conscious sedation using i.v propofol 2 mg/kg/h and remifentanil <0.2 g/kg/min, adjusted to the specific needs of each patient. Patients’ O2 Sat and expired CO_2_ levels were monitored by capnography.

An endocardial map of the right ventricular outflow tract was obtained on CARTO 3® using a mapping catheter (Pentaray™ NAV Eco, 2-6-2mm spacing, or SmartTouch™, Biosense Webster, California, USA), passed via the right femoral vein and areas of abnormal electrograms (definition below) were annotated if identified. CARTO-UNIVU was used in one of the centers for integrating electroanatomical mapping and live fluoroscopy. Electroanatomic data were collected either point-by-point or using an automated point collection facility (CONFIDENSE™ Continuous mapping) as per operator preference. The automated system relied on filters for each beat based on a series of pre-assigned parameters. Criteria for including points were (1) a cycle-length stability within a 5% range of the R-R interval; (2) an electrode position stability within 2 mm; (3) gating to end expiration of the respiratory cycle; (4) a contact force above a minimum of 5g (to avoid sites with insufficient contact). To ensure adequate surface contact, a tissue proximity filter which only collected points in proximity to the endocardial/epicardial surface based on an impedance measurement algorithm of either dipole was utilised.

Abnormal electrograms were defined as those having (1) low voltage (<1 mV); (2) split electrograms or fractionated electrograms with multiple potentials with 2 distinct components, with 20 ms isoelectric segments between peaks of individual components; and (3) wide duration (>80 ms) or late potentials, with distinct potentials extending beyond the end of the QRS complex, as suggested by Nademanee et al [5].

Pericardial access was obtained through percutaneous subxiphoid puncture with a *Tuohy* needle (or pericardiocentesis needle) as described by Sosa et al [16, 17].

Once within the epicardial space, the catheter was manipulated to map the anterior wall and right ventricular outflow tract epicardial surface. The deflectable Agylis™ sheath (Abbott©, Illinois, USA) was used whenever required by the operator.

If patients presented with baseline type 1 Brugada pattern electroanatomical mapping was performed immediately, but if the patient was presenting with non-diagnostic Brugada ECG a drug-challenge was performed with flecainide (2mg/Kg over 10 minutes – Centre 1) or ajmaline (1mg/Kg over 10 minutes – Centre 2). Abnormal electrograms (defined above), were marked if present, and targeted for ablation. Area calculation software present on CARTO was used to accurately define areas of abnormal electrograms during sinus rhythm. All patients from Center 1 underwent coronary angiography prior to radiofrequency ablation to assess proximity to the coronary arteries (energy was applied only >5 mm from a coronary artery).

Ablation power was 40W and energy was delivered for periods of 30 seconds, or dragging, to all areas with abnormal electrograms (Table 2). Ablation points were marked manually, or using VISITAG™, as per operator preference. Standard settings were: 3-mm distance limit for at least 10 s and a minimum contact-force of 10 g over 50 % of the set time period with a target force-time integral ≥400 g*s, but adjustment was performed throughout the case if requested by the operator.

After ablation, patients were remapped and more applications were delivered in case more areas of interest requiring ablation were identified. The procedure end-points were absence of the type-1 pattern and/or ablation of all electrogram targets. If type-1 pattern persisted after the operator ablated all the identified targets by conventional mapping the procedure was considered finished, but the endpoint of Brugada ECG-phenotype eradication was not achieved.

Electrophysiological study with ventricular stimulation up to 2 extrastimuli in the RV apex and drug challenge after ablation were performed whenever the operator felt appropriate.

### Procedural Results and Complications

We assessed the number of arrhythmic events (ventricular tachycardia or ventricular fibrillation requiring ICD intervention, or self-terminating) following the ablation procedure. These were recorded, through device interrogation when the patient came to clinic (once every 6 months for routine appointment, or more frequently in case of symptoms), or through remote monitoring.

A 12-lead ECG was repeated in 3-month intervals or if symptoms, to assess for the presence of relapsed type-1 Brugada ECG pattern.

Clinical outcomes were defined as: freedom from ventricular arrhythmia and type-1 ECG Brugada pattern relapse.

Procedural safety, complications occurring during the index admission or leading to rehospitalisation, was assessed and included acute and chronic coronary arterial damage, phrenic nerve palsy, phrenic nerve palsy, pericarditis, cardiac tamponade, ventricular pseudoaneurysm, ventricular-abdominal fistula, liver puncture, hepatic subcapsular hematoma, intra-abdominal bleed, death, and other procedure-associated causes of re-hospitalization.

### Ripple-Mapping Analysis

Ripple Mapping is a new feature in the CONFIDENSE^TM^ Module in the CARTO 3^®^ System. Each electrogram is displayed as a dynamic bar with its base at the mapping site and height related to the bipolar electrogram voltage at that time point in the annotation window. Both positive and negative deflections of the bipolar electrogram are represented by an outward movement of the bar from the cardiac shell. Point collection within a small area produces a ripple effect, with each bar changing according to the voltage change in a temporally accurate sequence, without the need of annotation. This enables simultaneous voltage and activation data on the same map displayed as sequential movement of dynamic bars preserving all of the low-amplitude deflections of multicomponent electrograms.

Ripple-Mapping analysis was performed offline after the ablation procedure. The activity propagation pattern on the recorded bipolar electrograms at each area was determined by Ripple Map. In Ripple Map, the threshold of bipolar voltages for display as a dynamic bar was set at 0.03–0.25 mV, which prevented baseline electrical noise from being displayed on the map.

In sum, during Ripple propagation, for every specific point in the map, all of the electrogram will be shown. Basically, instead of having a specific point of the electrogram (wavefront or peak signal) being represented in the propagation, and therefore being dependent on which part of the electrogram is annotated, Ripple shows what is happening with signal amplitude (represented by bars), in the point the electrogram was collected, throughout the propagation. This means that areas of fractionated potentials would only be represented by once during a normal propagation map, while using Ripple mapping the area will display a bar throughout the cardiac cycle whenever electrogram activity is present. Areas of long duration potentials, using the above mentioned criteria, were identified by visual inspection of Ripple propagation, and corresponded to areas where following the end of the QRS complex dynamic movement of ripple bars was still present for > 80ms. An independent investigator (RP) blinded to the location of ablation during the case marked these areas of prolonged ripple activity in a still frame (Panel C – Figures 1 to 4), and agreement regarding their location was obtained between 3 investigators (RP, PA, and FMC).

**Figure 1.**
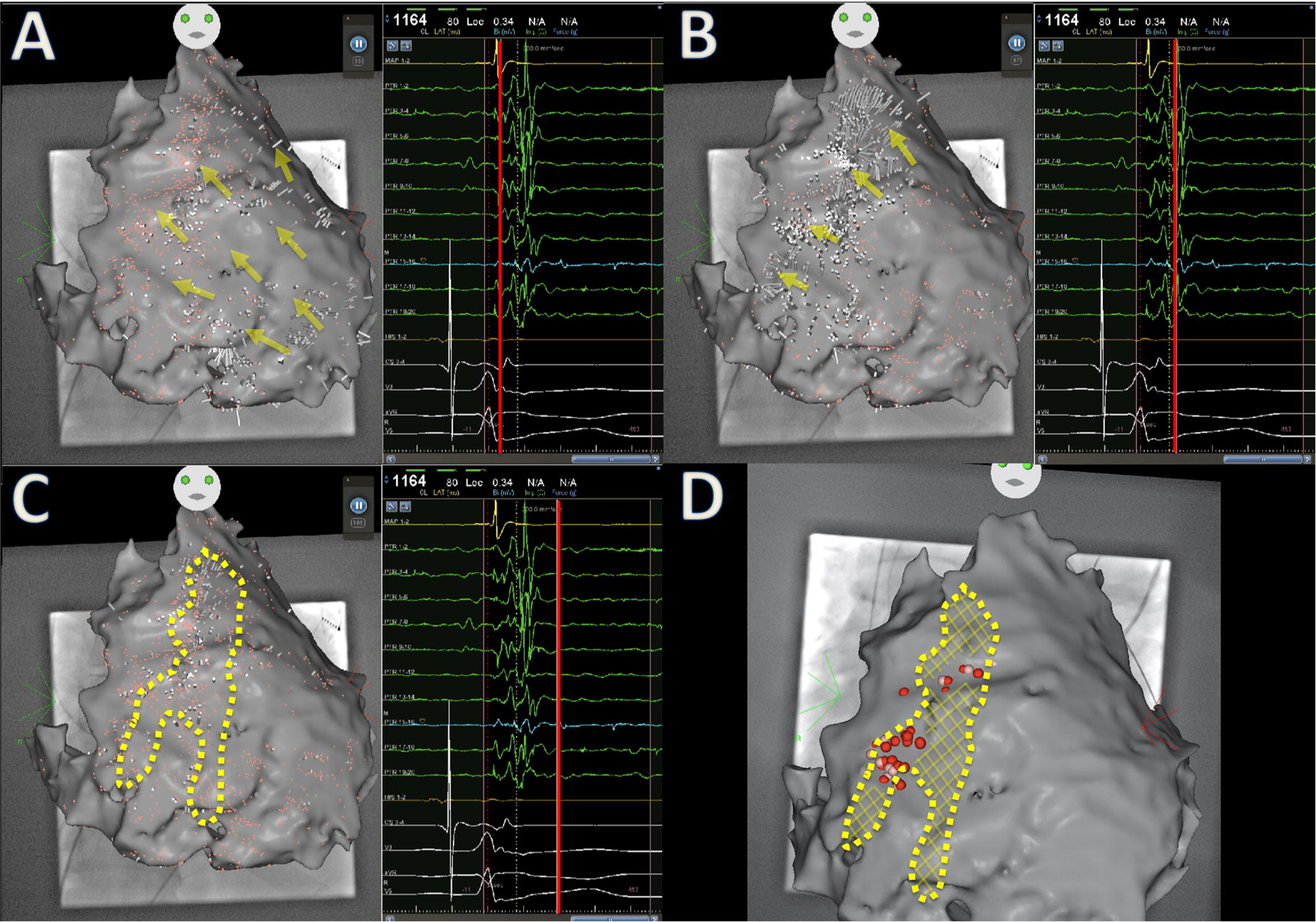
Electroanatomical map showing depolarization (direction or propagation illustrated by yellow arrows) of the epicardium of the RV anterior wall and outflow tract in Patient 1 (Panel A). Panel B shows areas of persistent Ripples during repolarization (highlighted by orange arrows). The same area is highlighted by a dotted yellow line (Panel C). Panel D shows how radiofrequency lesions (in red) correlated with the highlighted area, and areas of ripple activity where no ablation energy was delivered are highlighted in yellow.

Points in the culprit areas were reviewed to confirm their accuracy and rule out artefacts or baseline interference. Then, they were correlated with the location of ablation lesions (Panel D – Figures 1 to 4). If areas of prolonged ripple activity were not target during ablation, a yellow cross was marked in the area to illustrate that despite having abnormal electrograms no ablation was performed.

## Results

Four patients had ventricular fibrillation ablation (2 in each center) during the study period using CARTO. The Rhythmia™ mapping system was used for a fifth patient from Centre 1 who was therefore not included in the analysis.

### Patient characteristics

All patients were male, and age ranged from 32 to 54 years (Table 1). Ablation indications included recurrent ICD therapies in 3 patients, and patient preference in 1 subject. A type 1 Brugada syndrome ECG, either spontaneous or induced, was available for all patients.

### Procedural results

Long fractionated potentials were identified exclusively in the epicardium, and therefore all patients had exclusively epicardial ablation. As no fractionated potentials were located in the endocardium, we could not determine whether or not these are spatially related with the epicardial changes. Acute ECG normalization with disappearance of Brugada pattern occurred in all but one patient. No procedural complications were observed and all patients were discharged in the following morning. Additional details on the mapping and ablation procedure are available in Table 2.

### Ripple Map Analysis and Association with Extension of Radiofrequency Ablation

CARTO took up to only 30s to process and show the Ripple map. Analysis and interpretation took us less than 5 minutes for each patient, similar duration to what was described by *Linton et al.* [10]. Ripple analysis allowed the delineation of areas of interest in all four patients (Figures 1, 2, 3, and 4) (Video 1, Video 2, Video 3, Video 4). The 3 investigators were in agreement with regard to the location and extent of the defined areas. High-density mapping was performed in Patients 1 and 2, and this allowed a better and more detailed definition of the ripple map. Extension of the area of long duration potentials was wider in these 2 patients. Detection of long duration potentials was also possible in patients whose maps displayed less points, through repeated and careful analysis of ripple pattern after the QRS.

**Figure 2.**
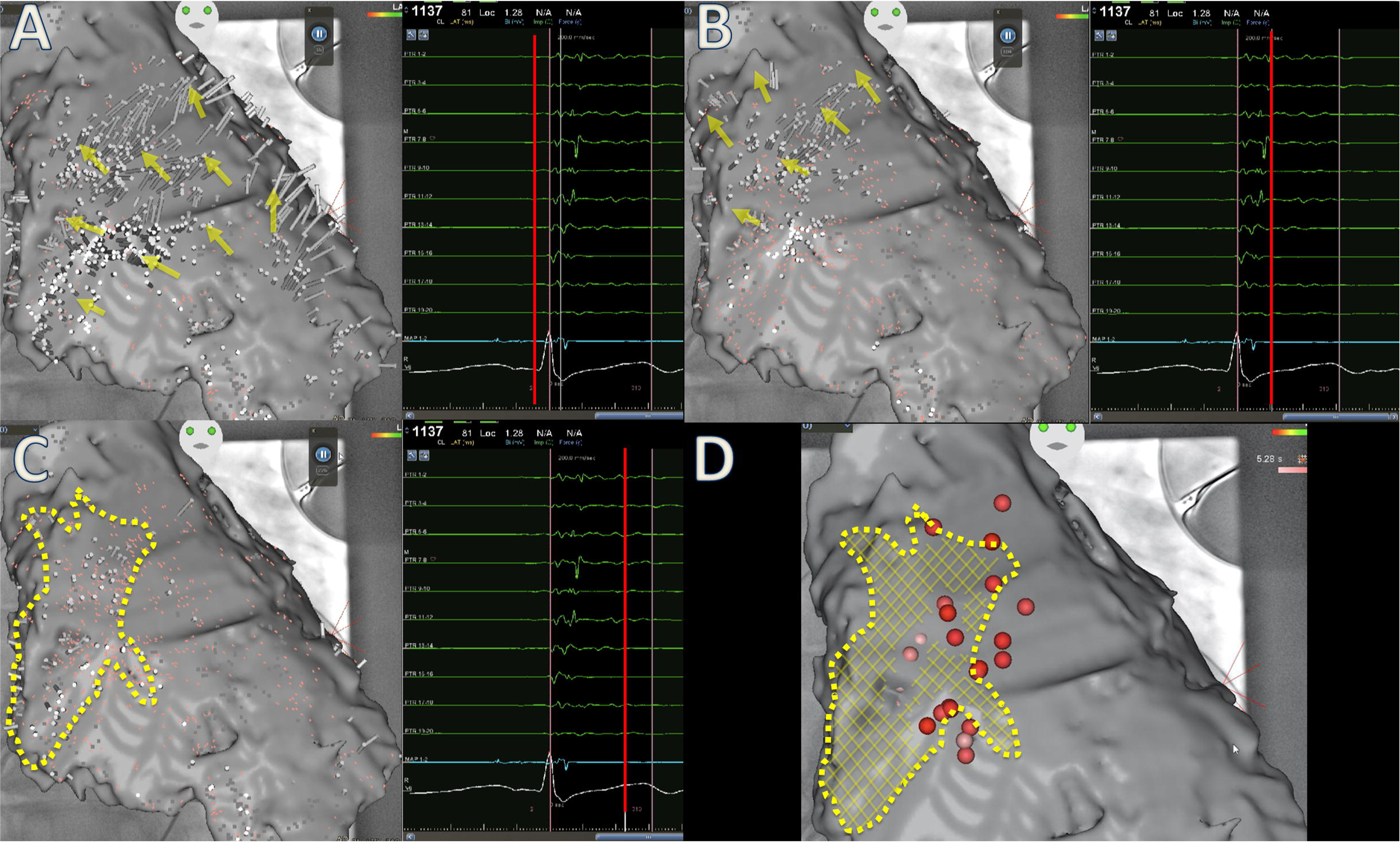
Electroanatomical map showing depolarization (direction or propagation illustrated by yellow arrows) of the epicardium of the RV anterior wall and outflow tract in Patient 2 (Panel A). Panel B shows areas of persistent Ripples during repolarization (highlighted by orange arrows). The same area is highlighted by a dotted yellow line (Panel C). Panel D shows how radiofrequency lesions (in red) correlated with the highlighted area, and areas of ripple activity where no ablation energy was delivered are highlighted in yellow.

**Figure 3.**
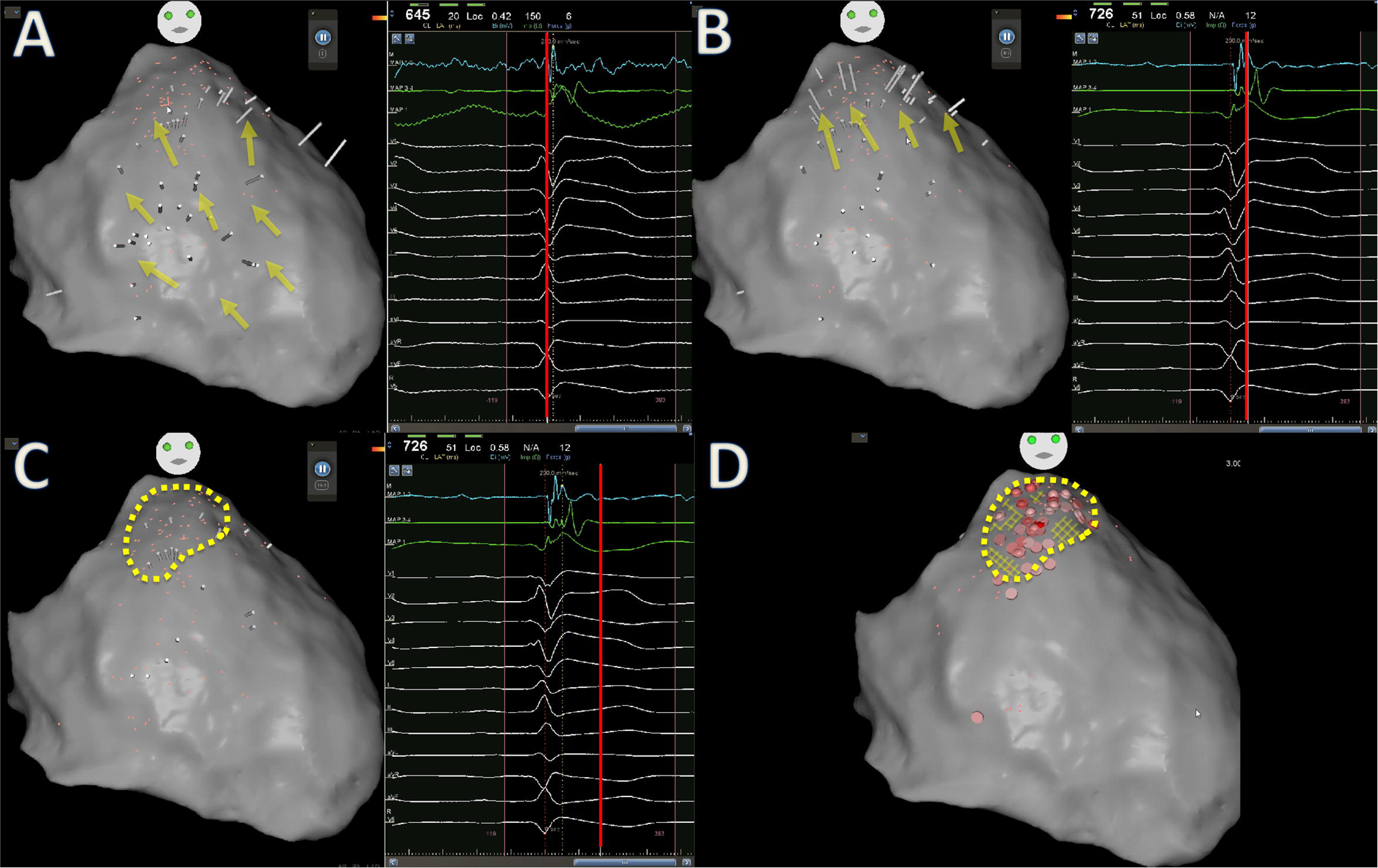
Electroanatomical map showing depolarization (direction or propagation illustrated by yellow arrows) of the epicardium of the RV anterior wall and outflow tract in Patient 3 (Panel A). Panel B shows areas of persistent Ripples during repolarization (highlighted by orange arrows). The same area is highlighted by a dotted yellow line (Panel C). Panel D shows how radiofrequency lesions (in red) correlate and cover nearly the whole area of interest.

**Figure 4.**
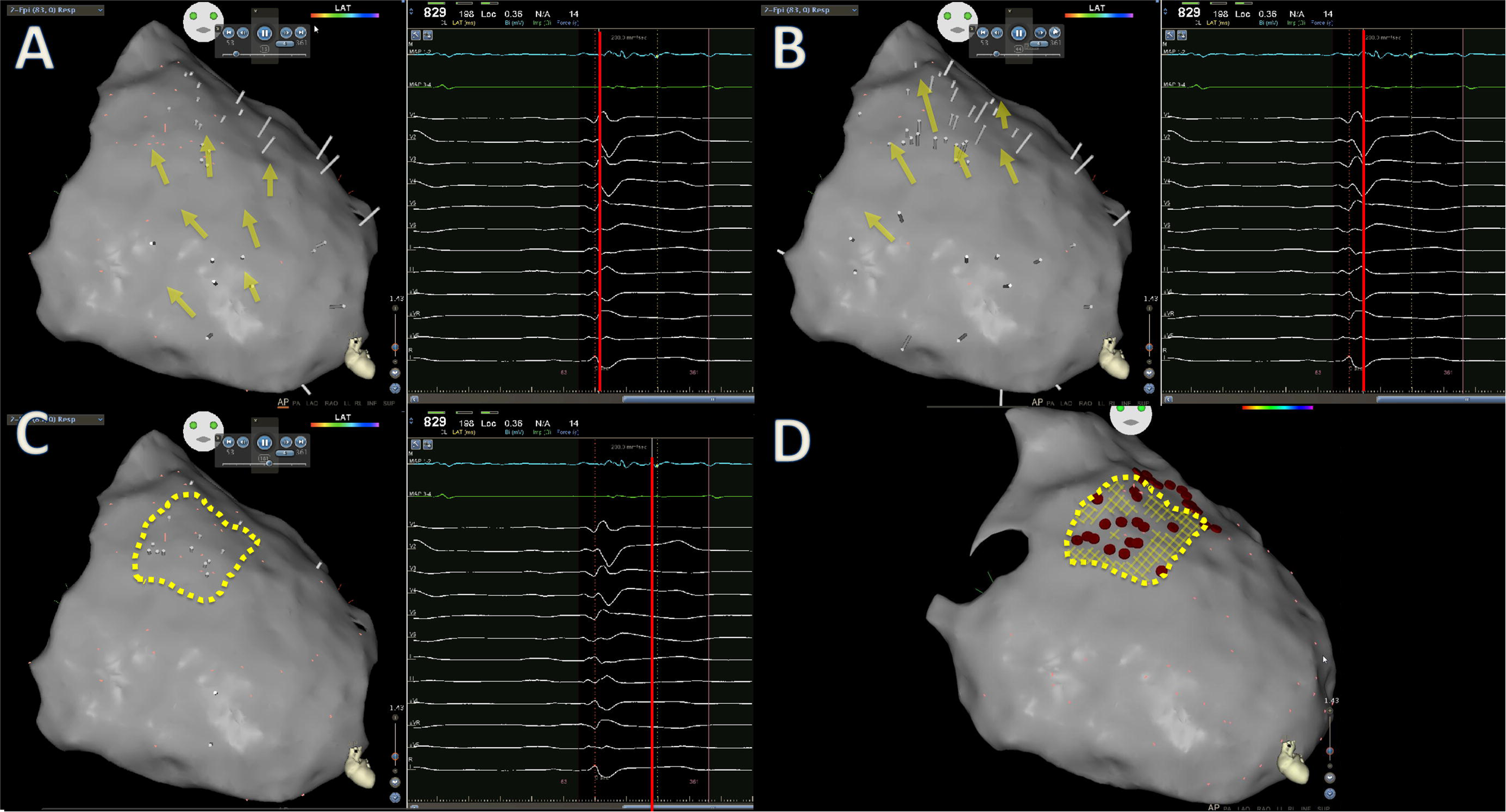
Electroanatomical map showing depolarization (direction or propagation illustrated by yellow arrows) of the epicardium of the RV anterior wall and outflow tract in Patient 4 (Panel A). Panel B shows areas of persistent Ripples during repolarization (highlighted by orange arrows). The same area is highlighted by a dotted yellow line (Panel C). Panel D shows how radiofrequency lesions (in red) correlate and cover nearly the whole area of interest.

Points marking the ablation area in patients 3 and 4 are a nearly perfect match to the areas identified by Ripple and standard mapping respectively. However, in patients 1 and 2, Ripple-mapping identified additional areas of long duration potentials which were not ablated. In patient 1 this area of non-ablated potentials appears account for > 75% of the region of interest, while in patient 2 it appears to correspond to <50%.

### Mid-term results

During a mean follow-up of 10.3±3.2 months all 3 patients with acute ECG-normalization presented with normal 12-lead ECGs during follow up, with no suggestion of relapse of Brugada syndrome pattern. However, Patient 1, whose resting 12-lead ECG had a persistent type-1 Brugada pattern, developed a further episode of ventricular fibrillation requiring defibrillator therapy, but all other patients were free from ventricular arrhythmia relapse (Table 3).

## Discussion

We have shown the potential use of Ripple-mapping to identify long-duration potentials in Brugada Syndrome patients referred for ablation in the setting of arrhythmic storm. Areas of multicomponent electrograms likely to be of interest for ablation were successfully identified on the basis of visual analysis of ripple maps and were found to correspond to the locations where ablation was performed in all 4 cases. Additionally, in 2 patients the analysis identified additional areas of abnormal electrograms. The clinical significance of these new multicomponent electrogram regions and the impact of ablation remains uncertain as our sample was small and not powered to assess this, but one of these patients with additional areas which were left unablated and corresponded to >75% of the whole region of interest presented with arrhythmia relapse. Had the operator been able to locate and ablate the remaining areas of fractionation (i.e. if Ripple had been made available to this operator) we can hypothesize that he might have succeeded in the procedure.

The mechanism by which substrate modification ablation prevents VT/VF in BrS remains uncertain. It is possible that by epicardial ablation, transmural dispersion of conduction-repolarisation is normalised to abolish the ST elevation on resting ECG and possibility of phase 2 re-entry [18]. Alternatively, by abolishing epicardial channels demarcated by fractionated electrograms as a result of conduction delay local re-entry and wavebreak degenerating into VF are prevented [19, 20]. Therefore, the exact size of the areas requiring ablation and endpoints of the procedure still remain to be determined as both mechanisms could be operative. Using automated mapping algorithms may ensure differences in procedural results arising from different levels of experience in signal interpretation are reduced and enable full eradication of abnormal signals if clinically indicated. Also, automated-collection of points with CONFIDENSE™ (in the 2 cases where we used CONFIDENSE™, >1,000 points were collected in 5 minutes) and subsequent analysis of Ripple without need of annotation may allow the identification of interesting multicomponent electrograms extending far away from the typical anterior right ventricular outflow tract region, to areas where operators would not be likely to spend less time carefully assessing electrograms, as in the lower part of the anterior right ventricle wall. However, this needs to be formally assessed in a prospective randomised trial. Nevertheless, through post-processing of data using Ripple-mapping we were able to quickly display and identify areas of abnormal electrograms which are known to be involved in the origin of ventricular arrhythmias in Brugada syndrome. Also, additional annotation or definition of window of interest was not required, which illustrates the simplicity and automaticity of the process. However, if operators require visual-help for assessing Ripple activity, it can be made even clearer if the post-QRS part of the sequence is played in loop repeatedly. CARTO also allows operators to visualize and manipulate catheters while the sequence plays, so that the areas can be better targeted and presence of fractionated potentials confirmed. Had we done a fractionation map for the same procedure [8] we would neither be able to see how the potentials display visually in real-time, nor get any information about sequence/propagation, and therefore we would be totally relying on the software analysis of electrograms and how well the algorithm performed.

As no interpolation is carried out by the Ripple-mapping algorithm, the more points are collected, the better the resolution of the map. This has been previously highlighted [21], and was clear in the two patients where we performed high-density mapping using the Penta-Ray catheter and CONFIDENSE™ to automatically collect/sample points. However, even in the two patients where point density was lower we were also able to identify the culprit areas.

Even though *Nademanee et al.* [6] and *Pappone et al.* [10] have performed quite extensive ablation of the RVOT in some of their patients, which could lead to the idea that purely anatomic guided-ablation of the whole epicardial anterior RVOT without mapping could be a treatment option, the total surface of ablated area varied widely among patients. This makes us believe that mapping is still crucial as we may not need to be very aggressive while ablating and may be able to treat *Brugada syndrome* patients with minor loss of RV function. Also, long-term data on the pro-arrhythmic potential of these epicardial lesions is still absent. Also, the fractionated potentials are not located exclusively in the anterior RVOT, and performing a Ripple map using CONFIDENSE™ can be a potential way of identifying additional targets in atypical locations.

Even though we did not identify areas of long-duration potentials in the endocardium we recommend that these should still be routinely looked searched for, as Haissaguerre et al [22], and also in our own experience using the Rhythmia™ mapping system [23] endocardial areas of interest can be found in some patients, but in that particular case the area of endocardial potentials had no anatomic relation with the epicardial changes observed in the same patients, as it was located to the posterior portion of the RV, close to the tricuspid annulus.

The electrophysiological basis for the Brugada phenotype is still a matter of debate, with two hypothesis receiving the widest support: nonuniform abbreviation of right ventricular epicardial action potentials (abnormal repolarization) vs. conduction delay in the RVOT (depolarization disorder) [24, 25]. Visual analysis of Ripple maps can provide some insight to what is happening in the RVOT, but probably will not provide the final and definitive answer to this important matter. We believe that ripples occurring shortly after the QRS may represent delayed conduction. Ripples occurring later after that (possibly 80ms or even longer after the QRS) may represent very delayed depolarization, or possibly repolarization heterogeneity, suggesting that both abnormal repolarization and delayed conduction may be involved.

Finally, this is the first publication showing the potential use of a commercially available automated mapping algorithm for catheter ablation of Brugada syndrome. Other groups [10] are in the process of developing and testing an algorithm created specifically for this purpose, but performance data are not yet available. Ripple-mapping currently depends on visual analysis. Possible improvements could include automatically displaying a map highlighting areas of longer ripple duration/density, measured above a certain cut-off point (to remove baseline artefact/or noise). These points could then be reviewed and confirmed before ablation.

This preliminary study has some limitations that need to be highlighted, namely the small sample size, short follow-up duration, and its retrospective nature. Ripple-mapping analysis was performed off-line after the ablation procedure, suggesting that analysis of ripples in high-density maps may be of interest to identify wider areas of long duration potentials. However, demonstration of feasibility of this technique in real-time in a live case is still lacking. The required extent of long-duration potential ablation, and the impact of ablation guided by ripple-guided analysis vs. conventional mapping should be assessed in future randomized controlled studies with sufficient power.

## Conclusions

This study shows the potential use of Ripple-mapping for identifying areas of long duration multicomponent electrograms which constitute a target for catheter ablation in patients with Brugada syndrome and recurrent ventricular arrhythmias and provide insight into its pathophysiological mechanism. At the moment, this technique allows the quick definition of an area of interest, which should then be validated by the operator before performing the ablation.

The impact of this approach on long-term outcomes compared to standard point-by-point analysis and operator-guided annotation should be tested in a randomized trial as a standardised analytical approach may optimise outcomes.

## Acknowledgments

none

